# Genomics-based annotations help unveil the molecular composition of edible plants

**DOI:** 10.1101/2022.01.24.477528

**Authors:** Shany Ofaim, Giulia Menichetti, Michael Sebek, Albert László Barabási

## Abstract

Given the important role food plays in health and well-being, the past decades have seen considerable experimental efforts dedicated to mapping the chemical composition of food ingredients. As the composition of raw food is genetically predetermined, here we ask, to what degree can we rely on genome-associated metabolic annotations to predict the chemical composition of natural ingredients? To that end, we developed an approach to unveil the chemical composition of 75 edible plants’ genomes, finding that genome-associated functional annotations substantially increase the number of compounds linked to specific plants. To account for the gap between the metabolic potential represented by functional annotations and the compounds likely to be accumulated and detected experimentally, we employed a predictive thermodynamic feasibility approach to systematically identify compounds that are likely to accumulate in plants. To quantify the accuracy of our predictions, we performed untargeted metabolomics on 13 plants, allowing us to experimentally confirm the detectability of the predicted compounds. For example, we observed 59 potentially novel compounds predicted and experimentally detected in corn. Our study shows that the application of genome-associated functional annotations can lead to an integrated metabologenomics platform capable of unveiling the chemical composition of edible plants and the biochemical pathways responsible for the observed compounds.

**Author Summary:** Nutrition and well-being take a central role in today’s high pace life, but how much do we really know about the food we eat? Here, we harness existing metabolic knowledge encrypted in staple food ingredients’ genome to help us explore the composition of raw edible plants. We first show the benefit and value of looking into genome-associated functional annotations on a wide scale. Next, we rely on new experimental data to develop a framework that helps us reveal new, potentially bioactive compounds in staple food ingredients. This has significance in, first, extending current food composition knowledge and second in discovering newly detected bioactive compounds, shedding light on the potential impacts of common food ingredients beyond their nutritional value as described in food labels. Finally, we show that staple foods that are already included in our daily diets might have the potential to contribute to our well-being.

## Background

“Make every bite count” is one of the main guidelines the U.S Departments of Agriculture (USDA) Dietary recommends for Americans (2020-2025)[1], reminding us of the multiple roles of the food we consume, and how it contributes to our well-being. Fruit and vegetables, serve as a source of energy and nutrients, modulating our health, and affecting disease [2–4]. Plants are complex organisms characterized by large genomes. For example, the corn genome (2,280 Mb) encodes 57,181 proteins, annotated to Gene Ontology (GO) categories, helping unveil the biological processes these proteins participate in and their potential molecular functions [5]. The genome can also be a strong predictor of phenotypic traits [6], like color, taste and aroma, known indicators of the nutritional value of a plant [7]. Indeed, color and pigments including carotenoids, betalains, and anthocyanins are well known for their bioactive properties [8]. Furthermore, the genetically predetermined polyphenols, alkaloids, carotenoids, and phytosterols have well-documented antioxidant and anti-inflammatory activities effecting multiple diseases, from cancer to diabetes or hypertension [9].

The composition of edible plants has been extensively mined in the context of bioactivity, focusing on antioxidants [10], anti-microbial potential as an alternative food preservatives [11], and most recently as holistic sources of bioactive compounds in work describing the use of the edible and non-edible parts of plant foods. For example, the non-edible parts of Bell pepper, like the seeds and leaves, were shown to be a valuable source of bioactive phytochemicals [12]. All these reports either rely on a collection of pre-existing work done on single plants or focus on a specific plant. To the best of our knowledge, the work presented here is the first systematic, large-scale approach developed for the exploration of edible plant composition.

Our current knowledge on food composition is limited to 150 nutrients catalogued by USDA, despite the fact that the true number of chemical compounds in food ranges from tens [13] to hundreds of thousands across all known plant species [14]. The bulk of our knowledge on the chemical composition of food comes from mass spectrometry and other low-throughput analytical methods and is compiled in repositories such as FooDB [15] and The Dictionary of Food Compounds [16] (DFC), cataloguing comprehensive information on the detected compounds, including both evidence-based and predicted annotations.

Our work is driven by the hypothesis that the full list of known, and yet unknown chemicals present in plants are encoded in the genome of the respective organism, encapsulating its metabolic capacities. Recent advancements in genomics have resulted in the emergence of extensive annotation efforts to decipher the genetic potential and the metabolic capacities of plants. For example, KEGG [17] links genes to their functional annotations such as enzymes, reactions, and chemical compounds and catalogues them in metabolic pathway maps, offering functional annotations for 7,254 organisms across the tree of life, out of which 56 are edible plants. Another contributor to plant genome-associated metabolic annotations is PlantCyc [18], a BioCyc [19] based platform adapted to annotate the functional diversity of plant genomes. Although the number of plant genomes sequenced, assembled and annotated has dramatically increased in recent years, the number of available genomes is relatively small (approx. 1,045 genomes in NCBI) and represents less than 0.16% of the known plant population [20]. As recent estimates indicate, approximately thirty domesticated plant species represent a significant portion of dietary diversity and three major grains (rice, wheat, and corn) contribute to more than half of the world’s caloric intake [21]. Thus, the exploration of existing genome-associated metabolic annotations of staple plant ingredients like corn, rice, soybean, potatoes, and other highly consumed edible plants offer a potentially good resource for describing their composition and hidden benefits.

Here, we explore to what degree existing genome-associated metabolic annotations can offer a valuable resource to deepen the knowledge of food composition. To do so, and focus on the edible parts of plants, we rely on metabologenomics [22–24], integrating genomics and metabolomics, used in the past to discover novel natural products [22,25]. To be specific, we develop a systematic metabologenomics approach, coupled with thermodynamic feasibility analysis, aiming to predict the composition of edible plants. We validate our predictions by comparing them to the existing chemical knowledge curated by food composition databases like FooDB, DFC, and USDA. We also collect new experimental data to explore the chemical composition of 13 plants. Our findings indicate that existing genome-associated metabolic annotations offer a predictive platform capable of systematically capturing the chemical composition of plant-based food, from fruits to vegetables, and allow us to predict and experimentally test the presence of novel compounds in plants.

## Results

### The existing knowledge on food composition

We collected data for 75 edible plants with published and metabolically annotated genomes from two well established databases: KEGG and PlantCyc. Our collection represents plants from 28 families including monocots and dicots (Figure 1a), covering major plant food groups: fruits (apple, banana and orange), grains (rice, corn and quinoa), vegetables (tomato, potato and spinach) and proteins (soy, chickpeas and pigeon pea).

**Figure 1.**
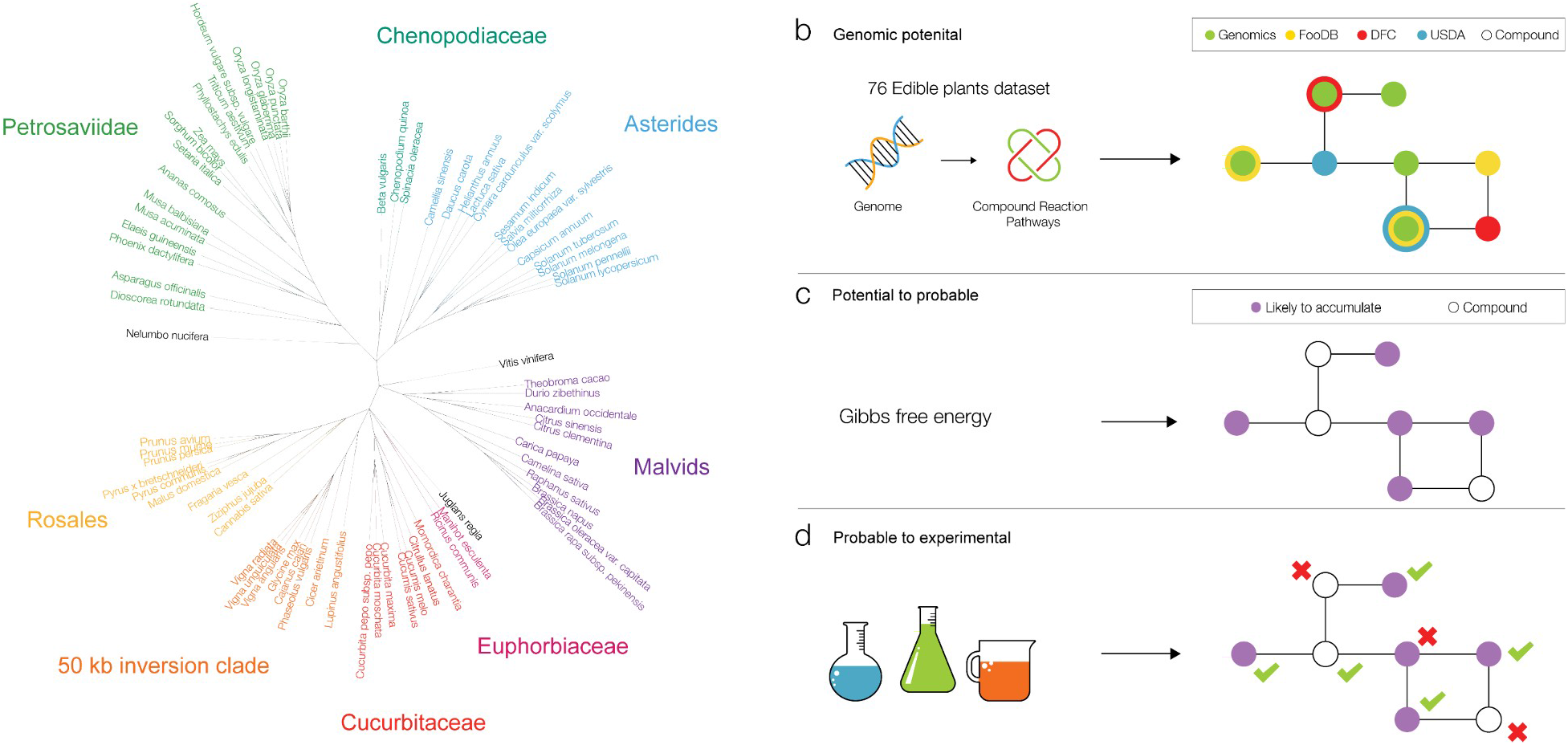
Plant phylogenetic diversity and schematic overview of genomics contribution to food composition knowledge. **(a)** Phylogenetic tree representing the 75 plants in our collection colored by plant order. **(b)** Collection and evaluation of genomics-based annotations to food composition knowledge. Functional annotations include compounds, reactions, and pathways. **(c)** Kinetics-based annotations help us to infer compounds likely to accumulate and hence experimentally detectable. **(d)** Validation of our kinetics-based approach against new metabolomics experiments, which detected compounds for 13 plants in our collection.

To estimate the currently available knowledge on the chemical composition of edible plants, we collected compound annotations from FooDB, DFC, and USDA, cataloguing 5,834, 5,151, and 140 compounds respectively, across all plants (Figure S1A). FooDB carries 723±452 compounds on average per plant (median: 850), a number that can be as low as three compounds for clementine (*Citrus clementina*) and as high as 2,181 compounds for tea (*Camellia sinensis*). DFC carries 83±121 compounds on average per plant (median: 33) with as low as one compound for vegetable marrow (*Cucurbita pepo subsp. pepo*) and red rice (*Oryza punctata*) and as high as 697 compounds for tea. Finally, USDA carries 88±46 compounds on average per plant (median: 110), with a single compound for false flax (*Camelina sativa*) and Chinese white pear (*Pyrus × bretschneideri*) and 128 compounds for apple (*Malus domestica*).

### Genome-associated metabolic annotations’ contribution to food composition

To estimate the contribution of available genome-associated metabolic annotations to the existing knowledge on food composition, we collected plant-related compounds from KEGG and PlantCyc, FooDB, USDA, and DFC. KEGG stores information about 7,245 organisms, out of which 546 are eukaryotes and 92 are plants, including 56 edible plants. PlantCyc is a plant-oriented database storing 126 genomes, out of which 58 represent edible plants. As all of databases detailed above are well-established and are of the highest quality of data, they provide annotations with sequence and literature evidence. Thus we consider the collection of plant-associated annotations as a comprehensive representation of the existing metabolic knowledge in the plant genome. These databases overlap and complement each other (39 plants overlap), covering 75 metabolically annotated plant genomes when combined. Overall, KEGG and PlantCyc contributed 1,201 and 3,737 new unique compounds respectively, adding a total of 5,224 new compounds (unique and common) to the composition of plants in our catalogue (Figure S1B).

To illustrate the contribution of genome-associated metabolic annotations to edible plants (Figure 1b), we focused on corn (*Zea mays*), a highly consumed staple crop [26] worldwide and in the US. Additionally, corn is known to have nutritional and medicinal qualities used in natural medicine to alleviate numerous symptoms, including pain and rheumatism [27]. Existing knowledge for corn includes 1,221 compounds from FooDB, 311 compounds from DFC, and 127 compounds from USDA. Considering overlaps between all sources, this compiled to a total of 1,038 unique compounds. Next, we set out to explore the value of adding genome-based annotations to corn’s existing knowledge.

One contribution of genome-associated metabolic annotations is the metabolic context of these compounds, carried by a network of pathways. Some known pathways are only partially annotated even after considering multiple databases. For example, in Monoterpenoid biosynthesis in corn (Fig 2B) six out of nine compounds are annotated in both databases. Of the three remaining compounds, (R)-Ipsdienol is known to be present in food but was not annotated to any plant in our collection. The two remaining compounds, Ipsdienon, a product of the reaction catalyzed by EC 1.1.1.386 directly from (R)-Ipsdienol, and (6E)-8-Oxolinalool, a product of the reaction catalyzed by EC 1.14.14.84, are currently documented in food but not in corn (white circles, Figure 2b). To strengthen the stringency of our work, the compounds, whose presence is documented in food but not known to be associated with corn, are not included in the plant’s catalogue.

**Figure 2.**
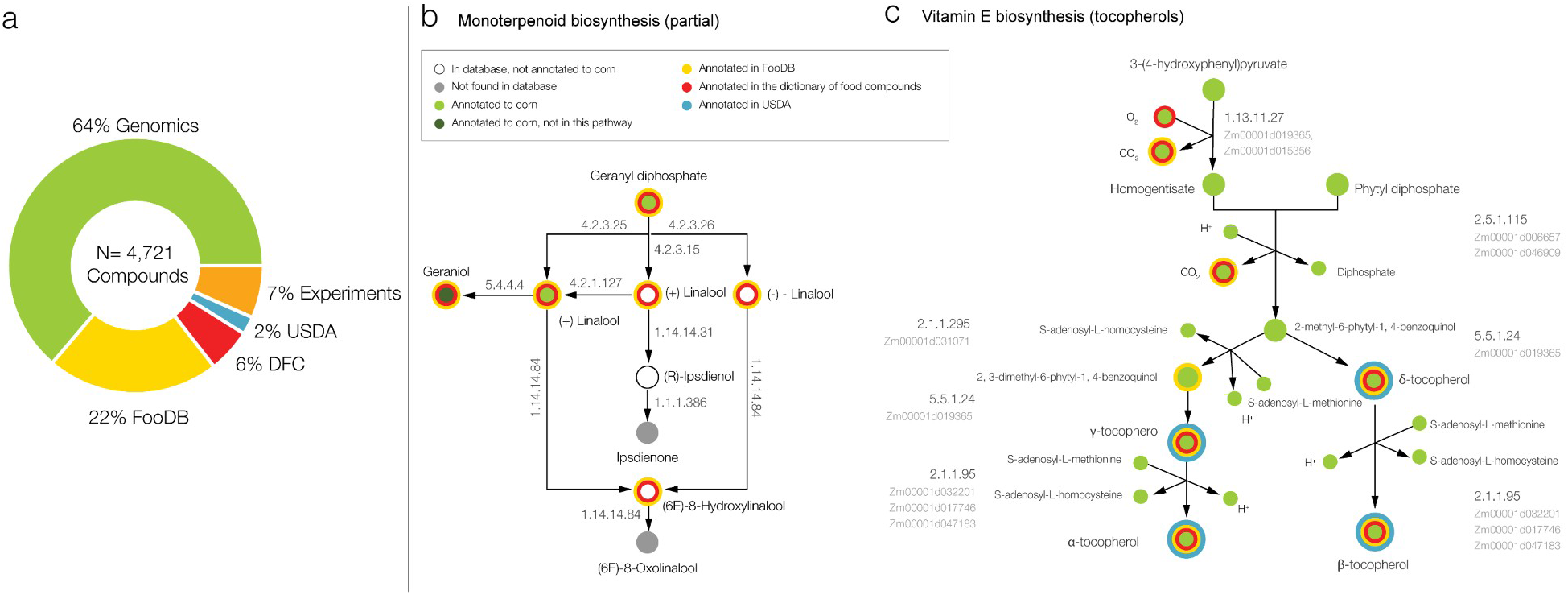
Genomics-based annotations boost corn composition knowledge. **(a)** Database annotations for corn, indicating that some compounds are annotated in multiple databases. The total number (N=4,721) represents the number of unique compounds for corn after the addition of genomics-based annotations. Genomics databases are represented by KEGG and PlantCyc. Other food related databases used in this work are DFC (the Dictionary of Food Compounds), FooDB, and USDA. Finally, experiments denote the set of compounds collected by metabolomics experiments reported here for corn. **(b)** A partial adaptation of the Monoterpenoid biosynthesis pathway in corn (KEGG) showing annotation availability, overlap and gaps in the coverage of different databases and genomics-based annotations. The different colors denote annotation sources as single or multiple concentric circles. **(c)** Vitamin E biosynthesis (tocopherols) pathway in corn (PlantCyc). Circles denote compounds and edges denote reactions. The different colors denote annotation sources as single or multiple concentric circles.

Consider another example, the tocopherol biosynthesis pathway. Tocopherols are an important class of compounds for health and nutrition [28] (α-tocopherol better known as vitamin E). Figure 2c shows the tocopherol biosynthesis pathway in corn and the delineated contribution of each database to its compounds. We find that databases such as FooDB and DFC annotate the lower half of the pathway, capturing products such as vitamin E and its derivatives. In contrast, genomics-based annotations offer a full pathway annotation, adding 4 new intermediates: 3-(4-hydroxyphenyl)pyruvate, homogenistate, phytyl diphosphate, and 2-methyl-6-phytyl-1,4-benzoquinol and 2 new cofactors: S-adenosyl-L-homocysteine and S-adenosyl-L-methionine, shedding light on the metabolic processes leading to the production of vitamin E in corn.

Taken together, existing genome-associated metabolic annotations added 3,021 compounds to the list of chemical compounds potentially present in corn, increasing its compound library by 64% (Figure 2a). Across our collection, genome-associated metabolic annotations increased the number of compounds by 2,363±728 chemicals on average per plant, an increase of 75%±15%. After this increase, our database documents 3,239±1,054 compounds per plant. We find that some plants are well annotated in FooDB, DFC, and USDA, while others are poorly annotated (Figure 3). Tea (*Camellia sinensis*) contains the highest number of FooDB, DFC, and USDA annotations (2,547 compounds out of a total of 4,379) showing an increase of 42% in new compounds. Some varieties of rice (*Oryza glaberrima*, *Oryza longistaminata*, *Oryza barthii*), white yam (*Dioscorea rotundata*), wild tomato (*Solanum pennellii*) and woodland strawberry (*Fragaria vesca*) are not catalogued by FooDB, DFC, or USDA, hence for these plants genome-associated metabolic annotations contributed 100% of the compounds. Other plants like clementine (*Citrus clementina*), lotus (*Nelumbo nucifera*) and African oil palm (*Elaeis guineensis*) are poorly annotated (<250 compounds) in FooDB, DFC, and USDA, hence the addition of genome-associated metabolic annotations increased our knowledge about their chemical composition by more than 85%.

**Figure 3.**
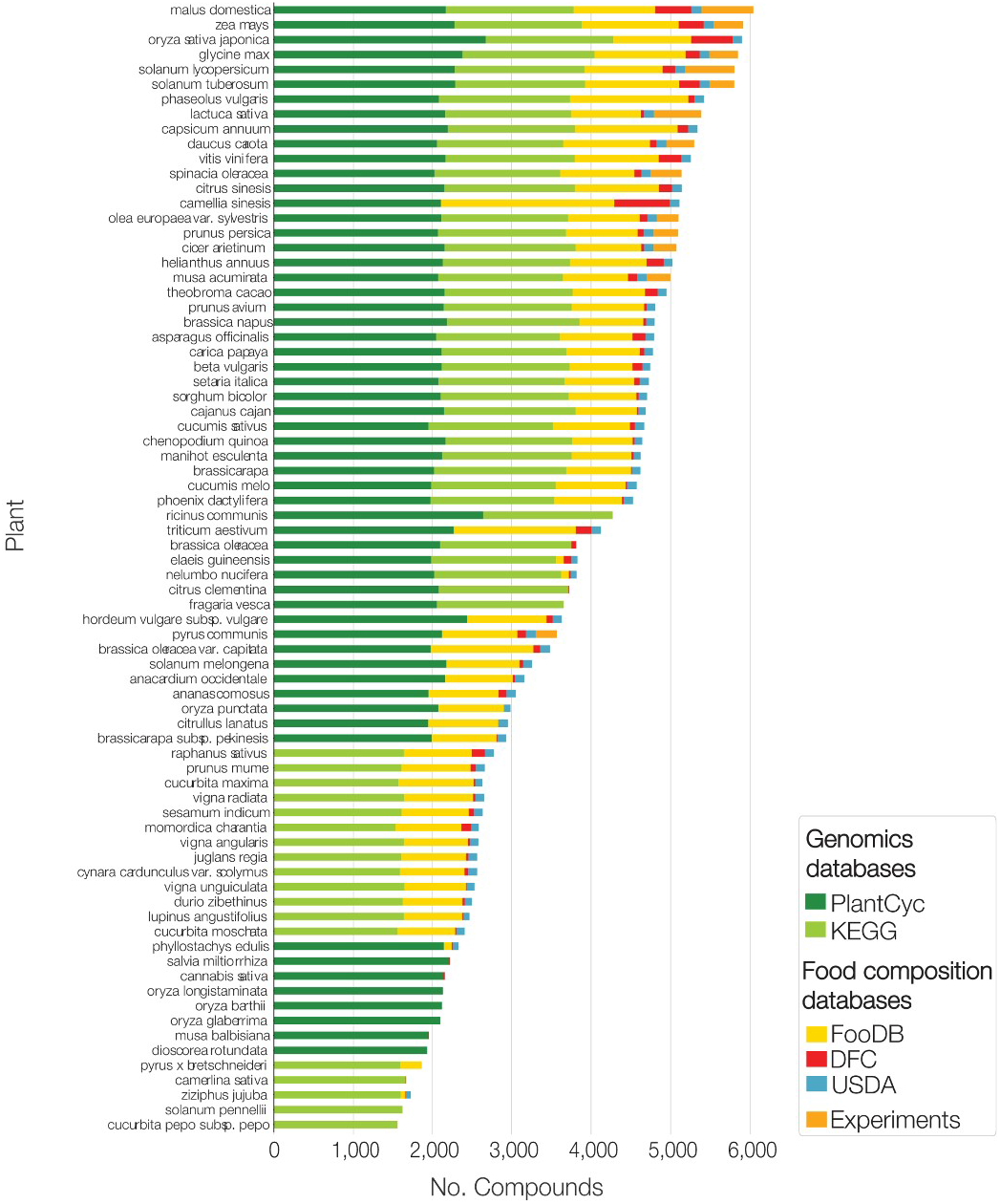
Contribution of the different sources of food composition across the entire edible plant catalogue. Genomics-based annotation are presented in two shades of green and include the KEGG (light green) and PlantCyc (dark green). Food composition databases include FooDB (yellow), DFC (red), and USDA (blue). Compounds detected in metabolomics experiments are shown in orange. The stacked bars represent here convey the proportions of annotations available from each of the databases/experiments. Annotations from each data source might overlap.

### Pathway enrichment analysis

Genome-associated metabolic annotations do not only increase our knowledge about the chemical composition of plants but also help us unveil the network of pathways responsible for the production of the newly predicted chemical compounds. Indeed, metabolic pathway mapping allows for a better understanding of the metabolic mechanisms responsible for the synthesis and modulation of natural products and offer a knowledge base towards the prediction of currently undetected compounds.

Metabolic pathways are divided into two main classes: primary and secondary metabolism. Primary metabolism is the collection of pathways involved in growth, energy, and reproduction of a plant, while secondary metabolism captures all other functions [29], like flavonoid biosynthesis, xenobiotics metabolism or plant-hormone metabolism, to name only a few. For corn, we collected 789 pathways, of which 35% belong to primary, and 65% belong to secondary metabolism. We asked if certain pathways are represented more than others in our catalogue. To test for pathway bias, we performed a hypergeometric enrichment test capturing the chance for a set of compounds mapped to a pathway in a certain plant to exceed the expected overlap with the general reference pathway (p-value <0.05) (Figure S2). Although the use of the hypergeometric test is a common and well-established tool, it is not without limitations, including recurring tests on the same pathway and the use of pathways as singular units with no inter-pathway biological dependence. To account for recurring tests on the same pathway, we applied the Bonferroni correction to the p-values in this analysis. The choice of database was shown to strongly affect pathway enrichment tests [30]. To maximize the biological relevance of our analysis and compensate for the effects that might rise from database selection, including biological dependence between pathways, we integrated pathway information from two inherently different pathway schemes; KEGG, with larger, more generalized pathway maps, and PlantCyc, a BioCyc-based mapping scheme with smaller, more specific pathway maps. In corn, we found 479 enriched pathways spanning both primary (45%) and secondary metabolism (55%), indicating that our corn dataset is metabolically diverse.

We next asked about specialized metabolism occurring only in corn, scanning our plant collection for enriched pathways specific to it (Figure 4a). Corn-specific pathways include specialized metabolism like kauralexin and zealexin biosynthesis [31,32], maysin biosynthesis [33], bergamotene biosynthesis [34] and all-trans-farnesol biosynthesis [35]. Each of these natural products of the diterpenoid, volatile sesquiterpenes, and flavone families are produced by the plant to acquire resistance against biotic and abiotic stress, such as herbivore attack.

**Figure 4.**
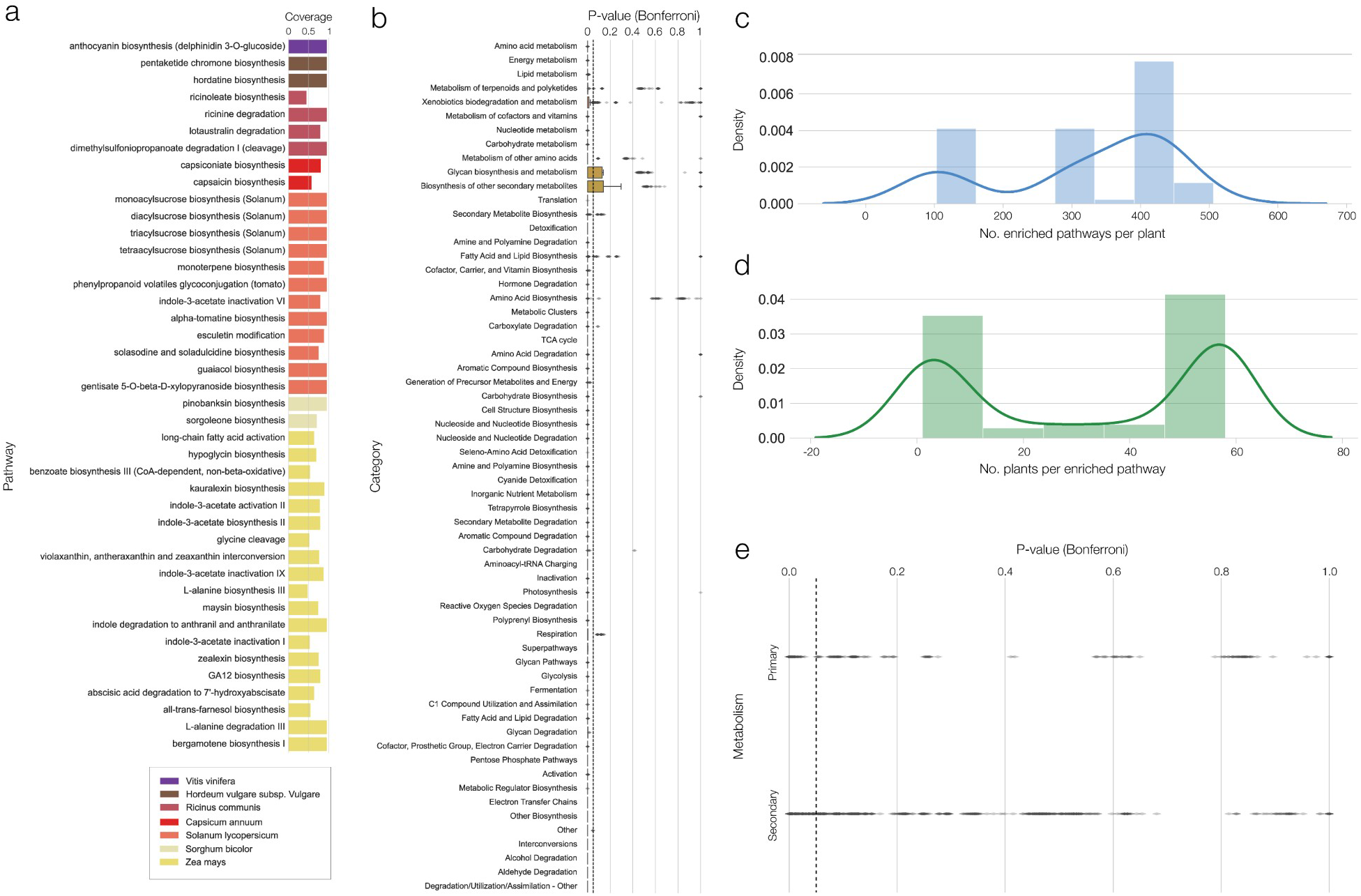
Pathway enrichment analysis. **(a)** Plant specific significantly enriched pathways (hypergeometric test, p<0.05, Bonferroni multiple testing correction). The coverage of each pathway is defined by the ratio of present compounds to the total number of compounds in the reference pathway. These include signature known pathways like Capsaicin biosynthesis in pepper (red bars), acyl sugars and alpha tomatine pathways in tomato (light red bars) and maysin and zealexin biosynthesis (yellow bars). **(b)** A boxplot of the p-values hypergeometric test, p<0.05, Bonferroni multiple testing correction) of pathway categories in corn. Medians found below the dashed line 0.05 enrichment line) represent the enriched pathways. We find that most pathways are enriched, indicating the diversity in our pathway coverage. **(c)** A density plot describing the distribution of enriched pathways per plant. We observed two peaks compatible with the two genomics databases included, KEGG (lower number of maps, each containing a larger number of reactions) and PlantCyc (larger number of maps, each containing a smaller number of reactions) **(d)** A density plot describing the distribution of the number of plants per enriched pathway. While most pathways are found in many plants, we observe several plant specific pathways described in detail in (a) and Figure S3. **e)** A boxplot of enriched pathways (hypergeometric test, p<0.05, Bonferroni multiple testing correction) classified as primary and secondary metabolism. Medians found below the dashed line (0.05 enrichment line) represent an enriched class.

Generalizing, the analysis performed for corn, we next classified pathways to primary/secondary in our entire collection. We identified 789 pathways across the full plant catalogue, 35% of which are related to primary and 65% to secondary metabolism, a fraction similar to the one observed for corn, offering evidence of diversity in our catalogue (Figure 4b, e). We find most pathways to be enriched (p-value <0.05), with the exception of pathways belonging to glycan biosynthesis and secondary biosynthesis metabolism. Other pathways belonging to secondary metabolism showed a large variation in p-values as multiple outliers were observed above the enrichment line (Figure 4b).

We found 762 enriched pathways across all plants (p-value<0.05), out of which 34% are related to primary and 66% are related to secondary metabolism. The distribution of the number of enriched pathways per plant shows two peaks (Figure 4c), corresponding to the two databases, KEGG and PlantCyc. The peaks capture the fact that KEGG has fewer, more complex, and generic pathway map representations, while PlantCyc has a larger number of smaller pathways. For example, KEGG represents the metabolism of alanine, aspartate, and glutamate in one map, while PlantCyc breaks down this process into 10 smaller pathways.

Different plants are known for their production of specialized compounds and natural products. Examples include the production of curcumin by turmeric, thymol by thyme, and vanillin by vanilla plants [36], prompting us to identify pathways enriched in single plants, pointing towards potential unique functionalities. Indeed, we observed a bimodal distribution of the number of plants per enriched pathway, one of the peaks being closer to 60 plants and another around 5 plants, indicating the presence of specialized plant metabolism in our catalogue (Figure 4c). Overall, we find that 30 out of 75 plants have at least one plant-specific enriched pathway (Figure S3), out of which we explored seven (Figure 4a). An example of a unique pathway is anthocyanin biosynthesis (delphinidin 3-O-glucoside) in grapes (*Vitis vinifera*), present in seeds and grape skins and used to differentiate between types of wine [37,38]. Other examples include: (1) hordatine biosynthesis in Barley (*Hordeum vulgare subsp. vulgare*). Hordatine, an antifungal compound, highly abundant in young barley shoots, reported to be a potential inhibitor of two main COVID-19 proteins, a protease (PDB ID: 7BQY) and a RNA polymerase (PDB ID: 7bV2) [39] and found in measurable quantities in different types of beer [40]. (2) Ricinoleate biosynthesis, found in castor bean (*Ricinus communis*) and the main constituent of castor oil, was shown to have antibacterial activities, and its elastic properties make it a candidate for packaging polymers with potential applications in biomedical and food technology [41]. (3) Capsaicin, a specialized component in pepper (*Capsicum annuum*) [42], was recently shown to have positive dietetic effects and beneficial antioxidant activity through association with the gut microbiome [43]. (4) Sorgoleone, an allelochemical exuded from sorghum (*Sorghum bicolor*) roots, is known to affect both microbial communities and neighboring plant growth [44]. Finally, we find a variety of acyl-sugar biosynthesis pathways in tomato (*Solanum lycopersicum*). Acyl-sugars are created in tomato trichomes and have commercial and medicinal uses [45]. Another unique pathway in tomato is phenylpropanoid volatiles glycoconjugation, a pathway describing specialized volatiles found in tomato fruits contributing to its signature smell [46].

In summary, publicly-available genome-associated metabolic annotations bring diverse metabolic information from both primary and secondary metabolism, offering evidence of new specialized compounds with bioactivity potential.

### Experimental confirmation of genome-associated metabolic annotations’ contribution to food knowledge

To experimentally test the capacity of existing plant genomics metabolic annotations to predict the presence of compounds in plants, we performed untargeted metabolomics experiments on 13 out of the 75 plants in our collection, resulting in a catalogue of 939 detected compounds (see Methods). All experiments were done on the edible parts of these plants. On average, 371±130 compounds were detected per plant, ranging from 264 in pear to 652 in apple, and 370 compounds for corn (39.4% of our experimental catalogue). These experimental results allow us to evaluate the accuracy of genome-associated metabolic annotations in the context of the edible parts of the plants analyzed. For this, we measured the overlap of the chemical structures between experimentally detected compounds and our known genomics-based compound collection, finding that genome-associated metabolic annotations significantly overlap with experiments (p-value= 0.018, SI section 1).

To be specific, we identify 59 compounds that are found only in corn genome-associated functional annotations and experimentally detected (Figure 5a). We clustered these new compounds based on their chemical structure and classified them into primary and secondary metabolism (Figure 5b, Figure S6), finding that the majority of compounds were attributed to primary metabolism (30) and might represent pooled intermediates and their possible fragments. Metabolite detection may also vary depending on the compound and spectra libraries used in identification. As primary compounds are better studied and annotated, secondary metabolites and their fragmentation products are less abundant in spectra libraries [47]. Overall, we observed well-characterized fractions of lipids, cofactors, sugar, and amino acid derivatives.

**Figure 5.**
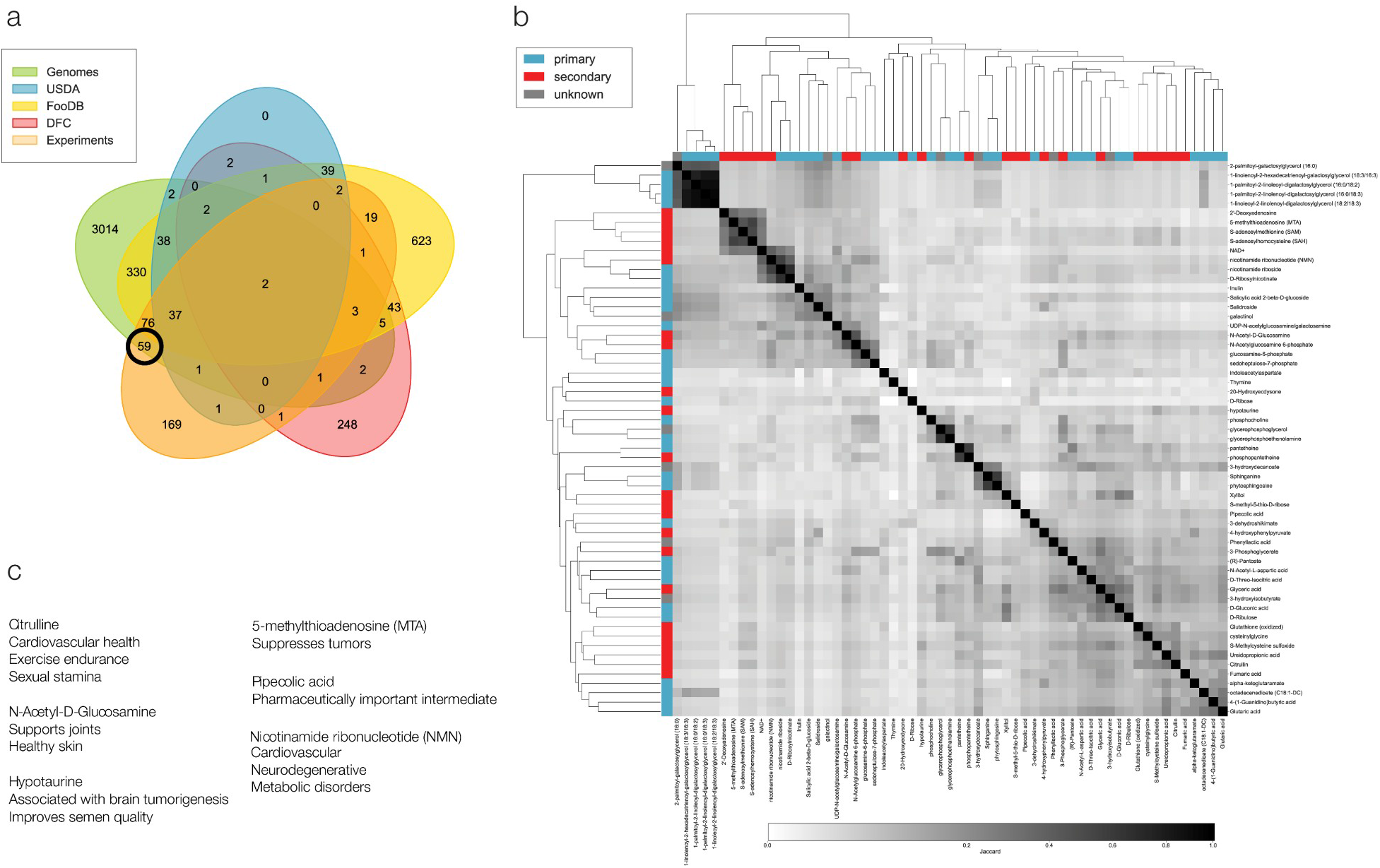
Bioactive compounds in corn unveiled by genomics. **(a)** Venn diagram comparing all sources of data contributing to the corn compound collection. The black circle marks the fraction of compounds unique to genomics-based annotations and metabolomics experiments reported here. **(b)** A structural similarity based clustermap of the 59 compounds highlighted by the black circle in panel (a). Compounds are classified to primary (blue) and secondary (red) metabolism according to the pathways they are part of. Greyscale denotes structural similarity as Jaccard distance, 0 being not similar and 1 being composed of the same bits in their vector representation. **(c)** The potential of corn as a nutritional influencer on well-being as learned with the addition of genome-based annotations, associated with antioxidant and anti-inflammatory activity promoting, heart, skin, and metabolic health.

We identified seven compounds present in corn that might potentially affect human well-being (Figure 5c). For example, Citrulline, a non-essential amino acid known to affect cardiovascular health and dilate blood vessels, and primarily found in watermelon and in smaller amounts in other fruits including a variety of corn species [48], is marketed as a dietary supplement for bodybuilders and athletes to improve exercise endurance. N-Acetyl-D-Glucosamine, previously detected in the shell that protects the first leaf of a corn shoot [49], is known to help support the joints and may help promote healthy skin [50]. Another compound detected is nicotinamide ribonucleotide (NMN), reported to be detected together with NAD+, a well-known cofactor, always present in the cell [51]. Well-being benefits related to this compounds stem from its effects on NAD+ content. This compound has been well studied and as a member of the vitamin B3 family [52] and is being used for the treatment of several cardiovascular, neurodegenerative, and metabolic disorders [53]. Other compounds detected in corn and unveiled by genomics-based annotations have potential cancer-related effects. For example, 5-methylthioadenosine (MTA), a sulfur-containing nucleoside, was recently reported as a tumor suppresser [54]. Another cancer-associated compound, hypotaurine, a sulfinic acid with antioxidant properties, derived from cysteine and an intermediate in taurine production, is one of the top-ranked metabolites for differentiating low and high-grade tumors [55] but was also shown to evoke a malignant phenotype in glioma, the most common primary brain malignancies in adults [56]. It is also marketed as a dietary supplement, together with taurine and L-carnitine, and associated with semen quality improvement [57]. Some other compounds observed in this subset were suggested to have both well-being and economic/industrial importance. For example, Pipecolic acid, a product of lysine metabolism, is an important regulator of immunity in plants and humans. In plants, it accumulates upon pathogen infection and is associated with systemic acquired resistance (SAR) [58]. Pipecolic acid is also an important intermediate of pharmaceutically and biologically derived compounds such as immunosuppressive agents and antibiotics [59,60]. The compound discussed above are known to have potential bioactivity for several applications. The potential effects related to medicinal, industrial and other applications, require further validation. Furthermore, rigorous quantitative analysis is needed to determine their contribution in the scope of quantitative consumption of raw or cooked corn or the concentrated form of a compound as a nutritional supplements.

Our experimental investigation also unveiled 22 compounds found only in FooDB, DFC, and USDA. These include: (1) 6-Methoxy-2(3H)-benzoxazolone (MBOA), a degradation product of the known bioactive compound 2,4-dihydroxy-7-methoxy-2H-1,4-benzoxazin-3(4H)-one (DIMBOA). DIMBOA is the main benzoxazinone synthesized in young corn tissues and accumulates in the cells. It is exuded by the roots and acts as a biocide against pests and as an attractant for soil bacteria. MBOA, the more stable form, is often detected in corn soils [61]. Interestingly, while DIMBOA is annotated to the corn genome, MBOA, its derivative is not. (2) Vanillic acid, a phenolic detected in corn grits [62]. (3) Trigonelline, an active alkaloid known to be found in corn and associated with antioxidant, anti-carcinogenic, anti-diabetic, and anti-hypercholesterolemia properties [63]. (4) Syringic acid, a phenolic compound found in fruits and vegetables including corn, is reported to have anti-oxidant, antimicrobial, anti-inflammatory, and antiendotoxic properties [64]. (5) Feruloylputrescine, a polyamine monoconjugate, previously detected in corn kernels [65].

Finally, to complete our survey of experimentally-detected compounds also found in our genomics-based annotations, we considered the remaining 12 out of 75 plants for which we have both experimental data and plant-genomics metabolic annotations. We find that the number of compounds added by genome-associated metabolic annotations is typically higher than those already catalogued by FooDB and DFC (49±15 and 26±14 compounds respectively, Figure S7). In other words, our analysis shows that genome-associated metabolic annotations significantly enhance existing knowledge of edible plants’ chemical composition, helping us uncover potentially novel bioactive compounds.

### Available genome-associated metabolic annotations annotations contribute to the feasibility of compound accumulation

The number of currently identified compounds detected by metabolomics is limited by instrumentation, standard libraries, and analysis pipelines. Recent work presented predictive approaches, applying existing knowledge as a basis for compiling lists of potential compounds that might be detectable by metabolomics experiments. One notable example of such an approach is MINEs [66], a tool that employs pathway and enzyme-based candidate rules to increase the amount and accuracy of compound annotations from raw mass-spectromtery data. Other approaches employ the existing knowledge together with thermodynamic principles to test the feasibility of metabolic pathways like glycolysis, explaining its activity under different conditions and constrains [67]. As thermodynamics rules and constrains are often used to provide insights on the pathway level, here, we hypothesize that the application of standard thermodynamic annotations, and specifically reaction Gibbs free energy values on the compound level can shed light on the likelihood of compounds to accumulate in different plants. Thus, we would be able to identify not only active metabolic routs but the compounds that might play a metabolic role or serve as important metabolic junctions in the plant cell. To that end, we set out to explore genome-associated metabolic annotations’ contribution to our ability to predict compounds that accumulate in a plant’s edible-parts. Indeed, chemicals that accumulate are more likely to be present in sufficient quantity to be experimentally detected or to enter the bloodstream, potentially modulating health. Indeed, transient compounds may be harder to detect.

To study the likelihood of a compound to accumulate, we used Gibbs free energy (ΔG) reaction values combined with genome-scale metabolic network topology. To establish thermodynamic feasibility, we determine the probability that a compound accumulates given all the reactions it takes part in a plant’s network context (Figure 6a). In other words, thermodynamic feasibility offers a method for discriminating compounds as likely to accumulate based on cumulative ΔG values of the reactions it is either a reactant or a product in. We collected ΔG values from modelSEED [68] and PlantCyc and calculated the cumulative ΔG value (score) for each compound. ModelSEED values are calculated as described in eQuilibrator [69], a well-established source for Gibbs free energy values. PlantCyc values are calculated according to the BioCyc schema of computing Gibbs free energy as described in [18]. If the compound acted as a reactant in a reaction, we assume that it is consumed, hence it is a transient compound, assigned a negative ΔG value. In contrast, if the compound is a product of the reaction, it is assigned a positive value. Compounds with positive scores are produced more than consumed, thus they are likely to accumulate. The set of available reactions was considered for each compound, out of this set, the reaction representing the largest absolute ΔG value may shift the balance towards either the consumption or production of that compound. This, in turn may determine the metabolic pathway activity and the creation of intermediates or byproducts that may or may not accumulate if they are not consumed by other thermodynamically feasible reactions. In corn, we scored 2,985 compounds (63% of total compounds), out of which 1,460 have positive ΔG values, i.e., are potentially expected to accumulate given the context of reactions they are involved in.

**Figure 6.**
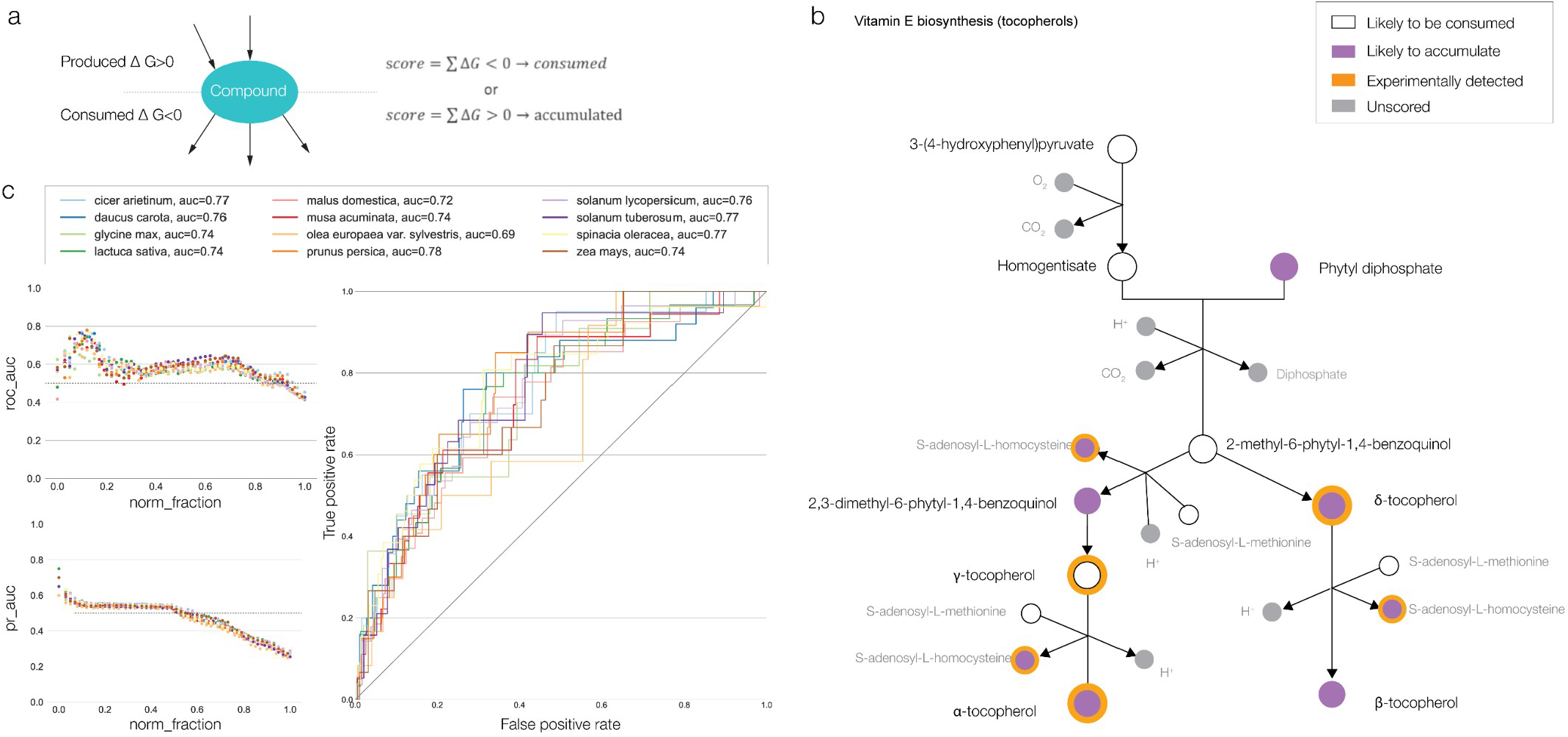
Application of kinetics-based annotations to predict likely to accumulate compounds. **(a)** Schematic description of the kinetics-based annotations approach. Gibbs free energy values, ΔG, were collected for each reaction in each plant and used to calculate the cumulative score of a compound. If a compound is a reactant in a reaction, it gets a negative ΔG value and if it is a product, it gets a positive ΔG value. **(b)** Vitamin E biosynthesis (tocopherols) pathway of corn (PlantCyc), describing the possible outcomes of our approach. Circles are compounds and edges are reactions. **(c)** Performance of our approach including (i) optimal threshold analysis on the top-ranking compounds predicted to accumulate, establishing the window of best performance denoted by the area under the receiver operator curve, AUC_ROC_ and the area under the precision recall curve, AUC_PR_ for each plant, and (ii) the optimal receiver operating curves for plants included in this analysis.

To illustrate our findings, we explored the compounds on the vitamin E biosynthesis pathway in corn (Figure 6b). We observed a potentially likely accumulation of pathway products α and β tocopherol and of the intermediates γ and δ tocopherol. Intermediates with the potential to accumulate are likely involved in more than one reaction outside the pathway. These reactions may have larger absolute ΔG values that can either lead to the rerouting of the compound to another metabolic pathway, creating a temporary hold on the production of certain compounds. For example, phytyl diphosphate is involved in 7 reactions and 5 pathways. The largest ΔG value measured for this set of reactions representing the largest ΔG value was annotated to the phytyl salvage pathway, describing the conversion of degraded chlorophyll to phytyl phosphate, contributing to its potentially high likelihood to accumulate. A compound could potentially accumulate if it is an intermediate appearing in two reactions with a large ΔG value difference in which the reaction with the larger ΔG value favors the production of the compound in question. For example, 2,3-dimethyl-6-phytyl-1,4-benzoquinol is involved in two reactions, both annotated to the vitamin E biosynthesis pathway, where the ΔG value for the reaction producing it (20.64 kcal/mol) is larger than the ΔG value for the reaction consuming it (-12.05 kcal/mol). The use of thermodynamic feasibility to predict potentially accumulating compounds that might be experimentally detected is not without its limitations. One example is pathway shifts due to ΔG value differentiation. These shifts are governed not only by thermodynamic feasibility but by various other cellular considerations such as genetic and metabolic regulation, and environmental conditions including nutrient availability, temperature, osmolarity, and pressure. To further estimate the power of our approach we condensed the compounds to compound families represented by the first block of their InchIKey. This allows for better comparison to metabolomics experimental results (see methods). Overall, 1,754±588 compound families were scored per plant, covering an average of 79±17% of its compounds. The fraction of scored compounds was as high as 96.5% of the total compounds in *Oryza longstaminata*, a species of rice, and as low as 50% in adzuki bean (*Vigna angularis*).

To estimate the predictive power of our approach, we first asked if kinetics-based annotations have increased the predictive power of our approach compared to total genome-associated metabolic annotations. We find that thermodynamics-based annotations show a more significant overlap with the experiments compared to genome-associated metabolic annotations (p-value=0.0018, SI section 1, Figure S4), and they are characterized by a high degree of structural similarity, significantly different from a random sample of the same size taken from genome-associated metabolic annotations (p-value<0.001, SI section 1).

Next, we used the experimentally detected compounds in the 13 plants for which we performed untargeted metabolomics, to estimate the performance of our approach. Similar to known machine learning methods, we look for compounds that are likely to accumulate, since our experimental data used as validation does not contain concentration information, we use compound score as means to establish the best performing threshold for solving for a binary classification (a compound might likely accumulate or not). After establishing the score threshold, all compounds with scores above it are considered as likely to accumulate. We then compare it to our experimentally detected compound catalogue as ground truth values (a binary classification denoting presence or absence). We calculated standard performance metrics, such as the true positive and false positive rates and the area under the receiver-operator curve (ROC), AUC_ROC_. In addition, we calculated the precision, recall, and F1 scores. Since our data may be imbalanced, we initially set the threshold of prediction to be larger than zero (positive values). We then performed a moving threshold analysis (see Methods) to determine the optimal threshold for best performance in each plant found in both our annotation and experimental catalogue (Table S1). Most AUC_ROC_ values were above the discrimination line (0.5), several representing acceptable discrimination (AUC_ROC_ values between 0.69 to 0.76) (Figure 6c).

Finally, we selected the top ranked 110 corn compound families with the best performance of our thermodynamics-based approach (AUC_ROC_=0.74). This list contained 15 experimentally detected compounds including AMP, S-adenosylhomocysteine (SAH), and 5-methylthioadenosine (MTA), which are cofactors maintained as constant pools in the cell. Other compounds such as succinate, Glycerol 3-phosphate, and 3-phospho-D-glycerate are key products of major energy producing pathways, such as carbon fixation by photosynthesis, glycolysis, and the TCA cycle. Interestingly, L-glutamate and D-Galacturonic acid were also detected. L-glutamate was previously reported as a key metabolite in corn, measured in large amounts in the endosperm [70,71]. D-Galacturonic acid is the main component of pectin, a polysaccharide naturally found in plant cell walls. Thus, both compounds are likely to accumulate and be detected in metabolomics measurements, supporting the predictive power of our approach. To further examine the possibility of detecting natural products in both our experiments and predictions, we compared our results to CoconutDB [72], an open collection of natural products. Indeed, we found that most of our predicted and experimentally detected compounds overlap with this dataset (Figure S5, Table S2). Surprisingly, CoconutDB contains many annotations for primary metabolites, including TCA cycle intermediates and simple sugars. Interestingly, the fraction of experimentally detected compounds that does not overlap with CoconutDB is comprised mostly of fatty acid derivatives. Indeed, the detection and annotation of lipids by mass spectrometry is limited. One of the limitations is the existence of thousands of structurally diverse lipids in large gradient of concentrations, some being below the experimental detection level. Another limitation in the detection and annotation of these compounds is ingrained in the differences between mammalian and plant lipids, as most of the existing lipid measurement technologies were developed for mammalian lipid analysis. Finally, lipid annotations are limited to well-studied and well-captured lipids. For example, the KEGG database contains 4,132 lipid annotations, representing only a small fraction of the expected diversity of hundreds of thousands of plant lipid species [73]. In many cases lipids are represented as a general backbone that can carry different residues (R groups). One example is the KEGG annotation of Glycerophosphoglycoglycerolipid (KEGG compound C06041), a generic compound with an unknown R group that still takes part in two pathways; (1) Teichoic acid biosynthesis and (2) Glycerolipid metabolism. Considering these and other limitations, the presented approach still captures a significant signal describing the compounds that can potentially accumulate in edible plants.

## Discussion

Here, we developed a systematic methodology to extract the contribution of publicly available genome-associated metabolic annotations to the molecular composition of foods. We found that genome-associated metabolic annotations not only boosted, in some cases by more than 85%, the number of compounds known to be present in a plant (in comparison to FooDB, DFC, and USDA databases) but also offered valuable mechanistic knowledge in the form of chemical structures and the metabolic pathways responsible for their production. Using multiple types of annotations (compounds, reactions, and pathways), we surveyed the contribution of genome-associated metabolic annotations in depth. These annotations, combined with experimentally detected compounds, were used to gain new insights into the chemical composition of edible plants, specifically corn.

The molecular composition of plants changes in time and in response to biotic and abiotic stresses such as environmental conditions, herbivore and pathogen attacks and hormonal signals, all governed by complex genetic and metabolic regulation. We therefore set out to explore the feasibility of a compound to accumulate, affecting the likelihood of being experimentally detectable. We find good discriminative power, supporting the large-scale use of thermodynamic annotations. As experimental data and thermodynamics annotations’ availability increase, they may lead to significant enhancement in the predictive power of metabologenomics.

While the advent of genome-associated metabolic annotations offers new insights into the composition of edible plants, it is not without limitations. Various biases that might arise from such data were explored throughout this work, representing only a few of the multiple factors that might affect our knowledge of food composition. One major limitation of a whole-genome holistic ‘metabolic-soup’ approach, specifically in complex organisms such as plants, is the spatial allocation of compounds. It is known that in plants, a compound can be formed in one tissue and accumulate in another. For example, the spatiotemporal distribution of amino acids, flavonoids, and more was shown to be differential across different tissues of citrus fruits with a larger fraction of flavonoids in the peel and an increased content of amino acids in the juice sacs [74]. Another challenge is whole-genome metabolic approaches is the integration of genetic and metabolic regulation. The use of transcriptomics in plants has been recently rising. As transcriptomics analysis advances, technologies such as single-cell transcriptomics and transcriptomics integrated with metabolomics are more widely used in plant biology [75,76]. One notable example is the integration of transcriptomics and metabolomics to study anthocyanin biosynthesis in cabernet sauvignon grape berries in response to growth condition changes [76]. A natural extension of transcriptomics is plant proteomics, describing the proteins that are encoded by activated or transcribed genes. Plant proteomics has also been shown to be valuable in agricultural and food applications [77,78]. The integration of transcriptomics and proteomics data into the approach developed here, may improve and augment the prediction of compounds that are likely to accumulate and be detected by mass spectrometry analysis in the future. Annotations of regulation of gene expression and metabolic feed-forward or back loops are limited. With the advancement of the collection, analysis, and standardized curation, these data will improve our predictions. Another major limitation is the availability of annotated plant genomes. One limitation is the ability of existing annotation tools to comprehensively annotate compounds that might be metabolized by enzymes encoded by orphan genes or genes with unknown functions. Recent work suggested such genes to contribute to a wide variety of biological functions and playing essential roles in plant phenotype [79]. As many annotation tools rely on homolog information to infer metabolic annotations (notable examples are plantiSMASH [80] and PlantCyc), these functions might be overlooked. Other metabolic annotation repositories like KEGG curate their annotations based on experimental and published work and therefore provide a more accurate collection. Still these data repositories rely heavily on published work that in turn might overlook orphan gene contributions. As plant orphan gene identification and study of function is rapidly advancing [81], improved annotations may soon be available for a more accurate investigation into the metabolic annotation of edible plants.

While the cost of sequencing dropped significantly, we find that the number of non-model plant genomes annotated remains limited and grows slowly. Other factors such as annotation quality and database standardization can also limit omics-based analysis. As shown here, two major databases, KEGG and PlantCyc, has introduced some redundancy in pathway mappings, partially rooted in the different pathway definitions used. As data continues to accumulate, standardization and mappings between the different data sources is key to deriving new insights.

The use of metabolomics to detect compounds in edible plants highly depends on the instrumentation used, standard libraries, and identification methods. Sample variation in the form of produce origin, growth conditions, and post-harvest treatments can also increase the variability and introduce environmental compounds that might be experimentally detected. For example, it is known that mass-spectrometers have poor detection of stereoisomers [82–84]. In our results, we apply the most stringent gold standard metabolomics annotation method, comparing the resulting spectra to known standards. To determine this approach’s comparability to the benchmark of massSpec annotation methodologies used for food metabolomics, we assessed the methodologies on the three most common experimental platforms (HILIC, RP, GC) and five food items (seaweed, Sichuan peppercorn, Miso, Pig’s Ear mushroom, Macaroni and cheese) and three annotation approaches: gold standard, spectra matching, and computational annotation. We find that the gold standard method tends to annotates up to 597 compounds, spectra library matching annotates up to 2,855 compounds, and the computational method annotates up to 5,225 compounds. We find that spectra library matching offers a 5-fold increase to gold standard and computation annotation provides an 8-fold increase. In our pilot the existing single-peak based tools annotated 2,426 peaks of the total 28,000 peaks combined over three experimental methods. Studies have shown that about 50% of the peaks in any MassSpec experiment are adducts [85–87], molecules formed by the addition of two or more molecules during sample preparation. This means that about 14,000 peaks correspond to individual compounds of which currently only 17.3% are annotated. Plants are known to produce around 200,000 compounds, including primary and secondary metabolites [88]. As new metabolomics annotation tools are being developed, with a focus on computational analysis, we expect more and more compounds to be annotated, increasing the discovery of a greater fraction of the vast diversity produced by plants. Even though considerable efforts have been made to establish spectral analysis pipelines, high-throughput metabolomics compound identification remains limited. Future improvements in metabolomics analysis and annotation could significantly enhance our knowledge of specialized metabolites and natural products in edible plants. Additionally, wider scale surveys including samples of produce from different points in the supply chain, starting from the field and ending in the grocery store will be useful to further study the accumulation patterns of potential bioactive compounds and their availability to the consumer.

Finally, the genome-associated metabolic annotations analyzed here greatly contribute to our existing knowledge of the composition of edible plants. Furthermore, this approach can be extended to be used for predicting microbial natural products and compound with bioactivity or industrial potential. Microbial genomes are smaller and easier to sequence and assemble. Moreover microbial species are better represented in the metabolic annotation databases and potentially may lead to more accurate predictions. The integration of genomics and metabolomics has been suggested as a promising combination towards the identification of promising compounds [89]. Here, we employ metabolomics of the edible parts of staple plant food ingredients and specifically corn to show the value this approach in the discovery of new compounds in specific plant tissues. Thus, this work offers a stepping stone towards a better understanding of food composition, offering insights based on fast-growing computational and experimental datasets.

## Methods

### Data collection

Compound annotations were collected from open-source (FooDB and USDA) and proprietary (DFC) databases. Genomics based compound, reaction and pathway annotations were collected from KEGG and PlantCyc. ΔG values were gathered from PlantCyc and modelSEED. All annotations were completed with SMILES, InChIKey, and molecular weight. Plant diversity was analyzed and visualized using the ETE3 toolkit [90] tree functions (ete3, version 3.1.2). All taxonomic data originated from NCBI. Overall, our catalogue includes 15,296, out of which 12,662 first block InChIKeys representing chemical families. To compare our catalogue and experimentally detected compounds (detailed in the next section), we collected first block InChikey compound annotations from CoconutDB (January 2022 release). CoconutDB identifiers were converted to BioCyc and KEGG identifiers for further functional analysis using CTS – The chemical translation service [91]. The database files are available on https://github.com/Barabasi-Lab/Plant-genomics.

### Mass spectrometry experiments

A selection of 13 produce items was purchased from two local grocery stores (Whole Foods Market and Stop & Shop): apple, banana, carrot, chickpea, corn, lettuce, olive, peach, pear, potato, spinach, soybean, and tomato. Produce available in grocery stores represent the product consumed by the populations. Indeed, this produce may be affected by post-harvest treatments and preservation methods that might introduce foreign non-plant produced compounds. To standardize and minimize these potentially foreign compounds, we treated the samples as they are eaten by the customer. For example, a common post-harvest treatment is applying a wax coating to fruit like apples, this protects the fruit from both mechanical and microbial damage. Removing the peel, removes these waxy compounds that otherwise would potentially be detected by mass spectrometry. Each produce item sample contained the combined material of six units (for example 6 apples) and was prepared in a humidity-controlled room with minimum light exposure. Sample preparation included peeling, chopping, freeze drying (-80°C for 24 hrs, Catalog No. 10-269-56B from LabConco/Fisher) and pulverizing into a fine homogenized powder (Kitchen Aid, 170W, Model No. BCG111OBO). All samples were prepared by the Giese lab (Northeastern University, Boston MA USA). The final samples were stored at -80 °C in vials containing 200 mg of powder and argon gas. The samples were shipped to two metabolomics centers (West Coast Metabolomics Center, UC Davis, CA USA and Metabolon, Morrisville, NC USA) for analysis on multiple platforms including UHPLC-CSH C18-HRMS-Orbitrap, UHPLC-BEH Amide-HRMS-Orbitrap, UHPLC-PFP-HRMS-Orbitrap, and Shimadzu LC and SelexION QTRAP MS and annotation (see SI section 2).

Experimental results were annotated by spectral matching or identified by a reference standard library. Results from both metabolomics centers were merged and standardized to InChIKeys (PubChem). Since metabolomics methodologies does not account for stereochemistry, only the first block of the InChIKey is used to compare two entries. Therefore, the resulting list is one of unique compound structures found in each food item. All mass spectromtery data is available in MetabolomicsWorkbench as ST002493 (for data acquired from UC Davis) and ST002492 (for data acquired from Metabolon).

### Statistics and enrichment tests

Enrichment analysis was performed with the hypergeometric distribution test (python 3.8, scipy [92] 1.6.3). Pathway enrichment test was performed per organism. To that end, we first collected all the compound annotations for the reference pathways from KEGG and PlantCyc, including all known annotations attributed to that pathway. We calculated the number of compounds found in each pathway in the organism and tested it for significance against the reference. To account for multiple testing on the same pathway, Bonferroni correction was applied to pathways’ p-values.

### Compound similarity across genomics, kinetics, and experiments

To investigate the degree of structural similarity and overlap between molecules retrieved by different techniques, we performed similarity search and clustering on a variety of molecular fingerprints (FP). We leveraged the python package rdkit [93] to standardize SMILES and InChIKeys associated with each compound annotation, using Morgan fingerprints (FP). To generate a Morgan FP, all substructures around the heavy atoms of a molecule within a defined radius are generated and assigned to unique identifiers. These identifiers are then hashed to a vector with a fixed length. The chemoinformatic community employs 1024- or 2048-bit vectors, populating them with fragments up to radius 1 or 2. For our analysis we increased the resolution up to 8192 bits and radius 3, to capture fragments of bigger size and reduce the potential bit collision.

Since the first block of the InChIKey represents several stereoisomers, we assign a bit vector to each first block as the union of all bit vectors representing the related isomers. We then compare the degree of structural similarity between any pair of compounds by computing the Jaccard similarity between binary vectors.

We assess structural similarity for a given set of *N* compounds, by calculating the *intrinsic dimension* of their Jaccard similarity matrix {*S_ij_* }, a function of the spectrum {*λ_i_* } of {*S_ij_* },

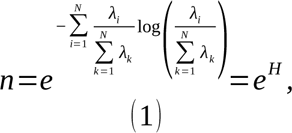

where the Shannon entropy *H* of the normalized spectrum is used to estimate the number of independent components synthetizing the same amount of structural similarity observed in the sample. A higher *n* indicates a more chemically diverse sample. We used *n* to assess how different the kinetics annotations are compared to a random sample from the genomics set (comparison with 1000 subsamples) (SI Section 1).

### Thermodynamic feasibility analysis

We collected available thermodynamic annotations for all the plants in our collection. Thermodynamic annotations in the form of ΔG values (Kcal/mol) were collected from ModelSEED[68] and PlantCyc [18].

As our validation set is based on metabolomics identified compounds, we used the first block of the InChIKey string as a compound identifier. First block identifiers collapse stereoisomer information, a known limitation of mass spectrometry. To calculate the cumulative score of ΔGs for a compound we collected all the reaction annotations in which it is involved. If the compound was acting as reactant in a reaction its ΔG value would be assigned a negative value and if it would be a product, a positive value. All the values were then summed to represent a likelihood to accumulate score where a positive value indicated a likely to accumulate compound and a negative value a likely to be consumed value. This approach was applied for each plant in our plant collection with corresponding experimental results. A schematic overview of the developed pipeline is described in Figure S7.

### Performance evaluation

To evaluate the performance of the thermodynamics feasibility approach, we used the first block of the InChIKey of our experimentally detected compound catalogue as a validation set and ground truth. For each plant we created a binary matrix representing the predicted score (calculated likely to accumulate value) and the experimental outcome (detected/not detected) using a cutoff threshold of a positive score (>0). We then created an ROC and precision-recall curve and calculated the area under the curves. To test for optimal performance, we first ranked our data according to the highest scores, implying better likelihood to accumulate and/or be detected experimentally. Next, we applied a moving threshold analysis to establish the cutoff threshold leading to optimal performance. Briefly, we systematically increased the portion of ranked data, calculating performance metrics for each step and identified the threshold leading to optimal performance by finding the maximum geometric mean of sensitivity and specificity, and updated performance metrics accordingly. All calculations were performed and plotted using the sklearn (version 0.24.1) and seaborn (version 0.11.1) python packages and Python 3.8).

## Acknowledgements

Experiments performed at the West Coast Metabolomics Center at University of California, Davis were led by Dr. Oliver Fiehn and Dr. Arpana Vaniya. Experiments at Metabolon were led by Nino Esile, Dr. Brian Ingram, and Nathan Testa. We thank Dr. John de la Parra, and Rebekah Carlson for preparing the food samples analyzed by UC Davis. We thank Dr. Roger Giese and Dr. Pushkar M. Kulkarni at the Department of Pharmaceutical Sciences, Northeastern University for preparing the food samples analyzed by Metabolon, Inc.

## Author contributions

**Shany Ofaim:** Data Curation, Formal Analysis, Software, Methodology, Validation, Visualization, Writing – Original Draft Preparation. **Giulia Menichetti:** Conceptualization, Formal analysis, Methodology, Software, Writing – Review & Editing. **Michael Sebek**: Data Curation, Investigation, Resources, Writing – Review & Editing. **Albert László Barabási:** Conceptualization, Funding Acquisition, Supervision, Writing – Review & Editing.

## Data availability

The datasets generated during and/or analyzed during the current study are available on https://github.com/Barabasi-Lab/Plant-genomics.

## Code availability

Code and scripts are available on https://github.com/Barabasi-Lab/Plant-genomics.

## List of Legends for Supporting Information Files

1. Supporting Information (PDF) – Contains all supporting text and supporting figures S1-S8.
2. Table S2 (CSV) – Contains supporting metabolomics data used in this work.

