## Supplementary material for "Genomics-based annotations help unveil the molecular composition of edible plants": Supplemetary material

### Supplementary information

Shany Ofaim[1], Giulia Menichetti [1,2], Michael Sebek [1] and Albert László Barabási [1,2,3]

[1]Network Science Institute and Department of Physics, Northeastern University, Boston, USA; [2]Department of Medicine, Brigham and Women's Hospital, Harvard Medical School, Boston, USA; [3]Department of Network and Data Science, Central European University, Budapest, Hungary.

### Section 1: Compound overlap and similarity across genomics, kinetics, and experiments

The chemical library compiled for this manuscript consists of $N_{library}=12{,}662$unique InChIKey first blocks, of which 885 were retrieved by the experiments, 4,698 were supported by genomic evidence, and 2,593 were confirmed by additional kinetic records (Figure S4A). We observe a significant overlap between genomic and experimental annotations, as quantified by an enrichment test with population size equal to $N_{library}$ (p-value=0.0181). Interestingly, the overlap between kinetic-based annotations and experiments shows an even higher degree of significance with p-value equal to 0.0018.

As explained in the Methods Section, once assigned a bit vector to each InChIKey first block, we proceeded in characterizing the structural properties of the kinetic-derived subset compared to the overall genomic annotations. First, we calculated the intrinsic dimension of both kinetic and genomic samples following Eq. (1). We find $n_{kinetics}=793.50<2{,}593$ and $n_{genomics}=1153.61<4{,}698$, indicating the presence of structural redundancy across the compounds in each set. Second, by sampling $N_{sample}=2{,}593$ InChIKey first blocks from the genomic pool for 1000 iterations, we characterized the statistical behavior of the intrinsic dimension for a random subset of genomic annotations, finding $n_{genomics}^{sample}=848.01\pm9.44$ (Figure S4B). Hence, kinetic annotations display a significantly higher degree of structural similarity compared to a random genomic sample of the same size, implying that they provide a different type of evidence.

### Section 2: Mass spectrometry experiments and validation

Samples were sent to two metabolomics centers, West Coast Metabolomics Center at University of California, Davis and Metabolon Inc., for testing on multiple platforms. UC Davis used a biphasic extraction protocol, Matyash, on a partition of the powder[1]. The organic top layer and the hydrophilic bottom layer are separated for analysis on different instrumentation. The organic layer was reconstituted with a methanol-toluene solution and separated into two aliquots, one for positive ionization and one for negative ionization on a reverse phase platform (UHPLC-CSH C18-HRMS-Orbitrap). The hydrophilic layer was reconstituted with an acetonitrile-water solution then separated into two aliquots, one for positive ionization and one for negative ionization on a HILIC platform (UHPLC-BEH Amide-HRMS-Orbitrap). Another partition of the powder was extracted with a methanol-water solution, reconstituted with an acetonitrile-water solution, and separated into two aliquots, one for positive ionization and one for negative ionization on a plant metabolite platform (UHPLC-PFP-HRMS-Orbitrap). Apple, basil, garlic, lettuce, strawberry, and tomato sample were analyzed on all three platforms and ionization modes. Metabolon used a biphasic extraction on a partition of the powder using standard methanol-water extraction. The organic layer was divided into three aliquots, two for positive ionization on a reverse phase platform (UHPLC-CSH C18-HRMS-Orbitrap) at two different parameters and one for negative ionization on the reverse phase platform. The hydrophilic layer was used on a HILIC platform (UHPLC-BEH Amide-HRMS-Orbitrap) in negative ionization mode. Another partition of the powder was extracted by Bligh-Dyer protocol and the organic layer was divided into two aliquots, one for positive ionization and one for negative ionization on a lipidomics platform (Shimadzu LC and SelexION QTRAP MS). 20 produce items (apple, banana, basil, black bean, carrot, chickpea, corn, garlic, lettuce, olive, onion, peach, pear, pepper, potato, spinach, soybean, strawberry) were analyzed on the first two platforms while only apple and basil were analyzed on the lipidomics platform.

The experimental methods described employ well studied and standardized technologies of metabolomics. For sample preparation, we used a derivation of a USDA protocol for food samples. Both service centers used traditional metabolomics methodologies, from common extraction protocols to standard instrumentation. The Bligh-Dyer extraction method dates back to 1959, meanwhile the Matyash extraction method is a derivatization of the Bligh-Dyer protocol. The methanol extractions used predate the biphasic extractions. In terms of instrumentation, the service centers use industry standard liquid chromatography to mass spectrometry instrumentation. Both service centers operate a hydrophilic method and a reverse phase method. The columns used are the most common for these two methods, the BEH amide for HILIC and CSH C18 for reverse phase. Over 90% of the services centers in the Metabolomics Association of North America (MANA) employ these methodologies[2]. MANA centers mostly note use in clinical and pharmaceutical samples in animal and human samples; however, the few plant focused centers also employ these methods in combination with some specific plant method. To detect plant metabolites, there two methods used, to use a specialized column such as the Pentafluorophenyl (PFP) as seen by UC Davis or more commonly a change in parameters of the reverse phase method as seen by Metabolon.

The composition results from both centers were compared to the two most cited databases for food composition, USDA and FooDB, on the foods in common (apple, basil, lettuce, strawberry, tomato) between the two service centers and the databases[3,4]. The selection of the food items from the databases were matched to the food items analyzed in experiments. For example, we analyzed the gala variety of apple with skin, so we chose the data for raw gala apple with skin within the food composition databases. Of the 150 nutrients within the USDA, it reports 53 compounds in apple, 53 in basil, 55 in lettuce, 57 in strawberry, and 56 in tomato. FooDB, an aggregation of literature-based sources, reports 975 experimentally confirmed compounds in apple, 918 in basil, 851 in lettuce, 859 in strawberry, and 1,037 in tomato. In total the experiments found 2,400 in apple, 2,602 in basil, 1,868 in lettuce, 1,849 in strawberry, and 1,879 in tomato. Apple and basil have the most compounds due to being analyzed on one more platform than the other foods. The results show that the experiments find vastly more compounds than both databases, and that the majority of the compounds have never been experimentally reported with only 27% of the compounds found in either the USDA or FooDB. About 51% of the compounds are in overlap between the two centers, showing there is more agreement between the centers than to previously reported compounds. In addition to the experimentally reported compounds of FooDB, there are also inferred compounds for each food which have no evidence. Of the inferred compounds the experiments were able to provide evidence for 200 compounds in apple, 263 in basil, 233 in lettuce, 255 in strawberry, and 250 in tomato. The total number of inferred confirmations is 302 compounds, demonstrating that the found inferred compounds are largely found in all food items.

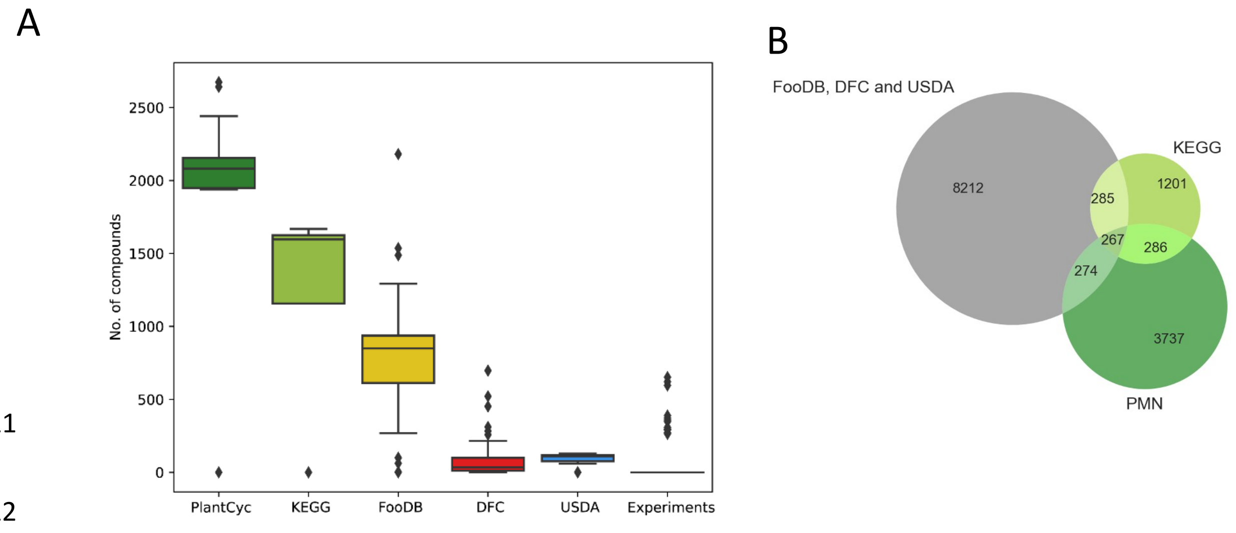


**Figure S 1 –** (A) Annotation contribution per plant for each database. (B) Complementation between existing knowledge and genomics-based compound annotations represented by Inchi Keys

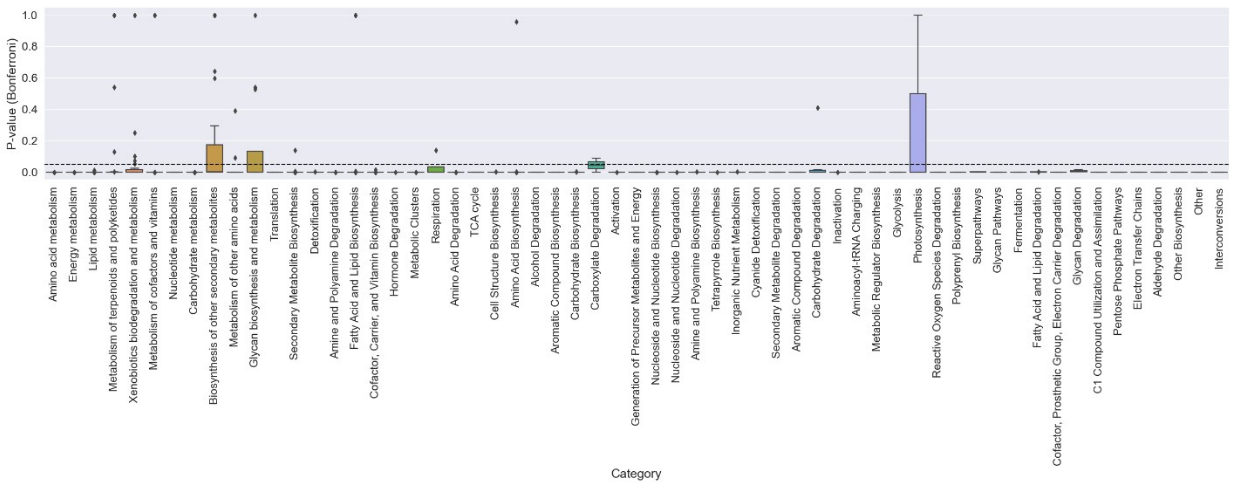


**Figure S2**- Analysis of corn pathways. Pathways found in corn were analyzed using the hypergeometric enrichment test (p-value < 0.05, Bonferroni) and binned into categories. P-values ranges across pathway categories are presented in a boxplot where medians found below the dashed line (0.05, the enrichment line) indicate significance (enrichment).

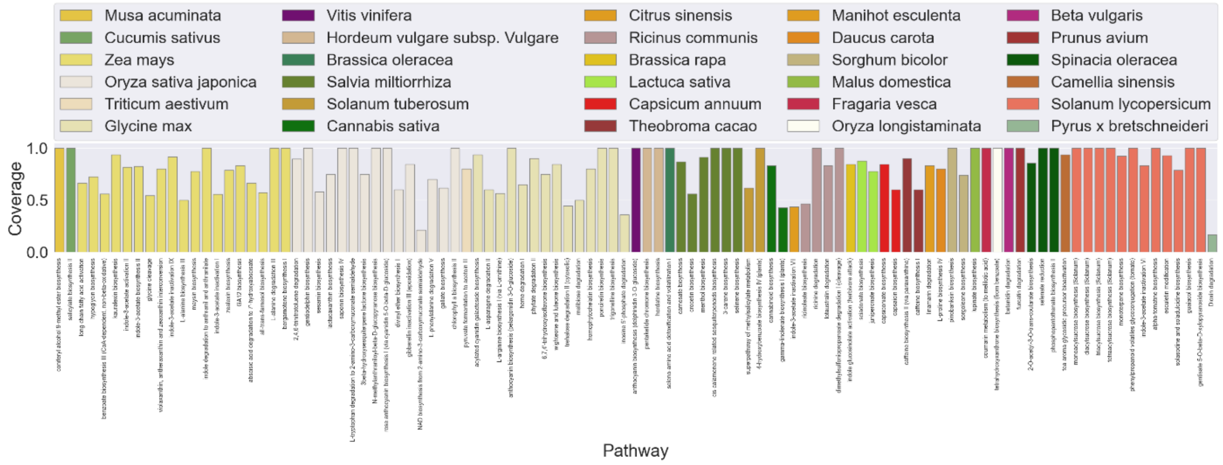


**Figure S3** - Significantly enriched pathways that are specific and unique to 30 out of the 75 plants in our collection. The coverage of a pathway is defined by the ratio of present compounds to the total number of compounds in the reference pathway.

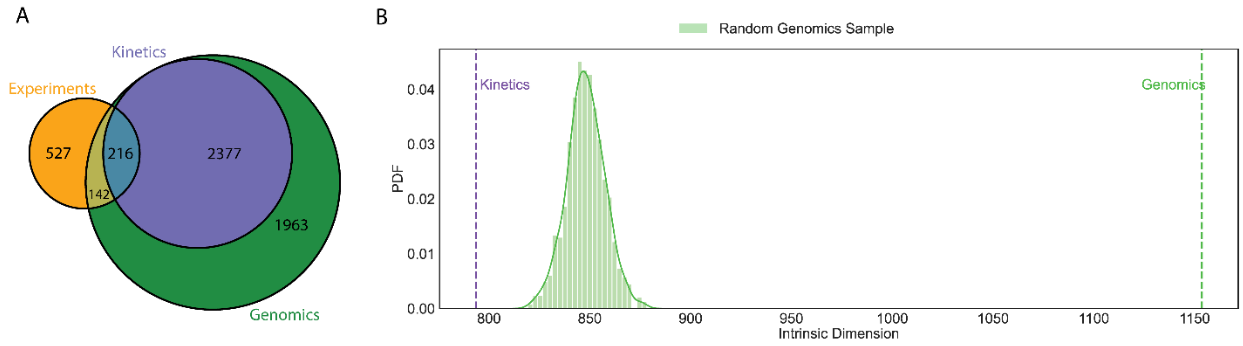


**Figure S4** – Compound redundancy and similarity across genomics, kinetics, and experiments. (A) Overlaps of InChiKey first blocks across genomics, kinetics and experiments. (B) Both kinetic and genomic annotations show some level of structural redundancy, as quantified by their intrinsic dimensions $n_{kinetics}=793.50$ and $n_{genomics}=1153.61$. Additionally, kinetic annotations are not compatible with a random sample of the genomic evidence, as over 1000 subsamples no one exhibits an intrinsic dimension lower than $n_{kinetics}$ ($n_{genomics}^{sample}=848.01\pm9.44$).

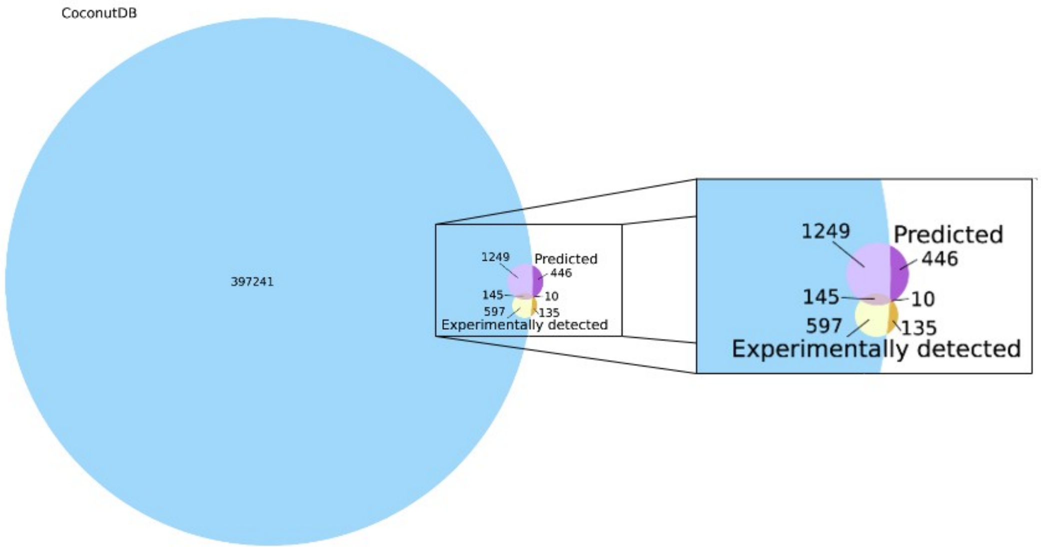


**Figure S5** – Comparison of our compounds that are predicted to accumulate (“Predicted”), compounds we experimentally detected (“Experimental”) and the compounds available from CoconutDB, a collection of natural products. The comparison was made by using the first part of the InchiKey.

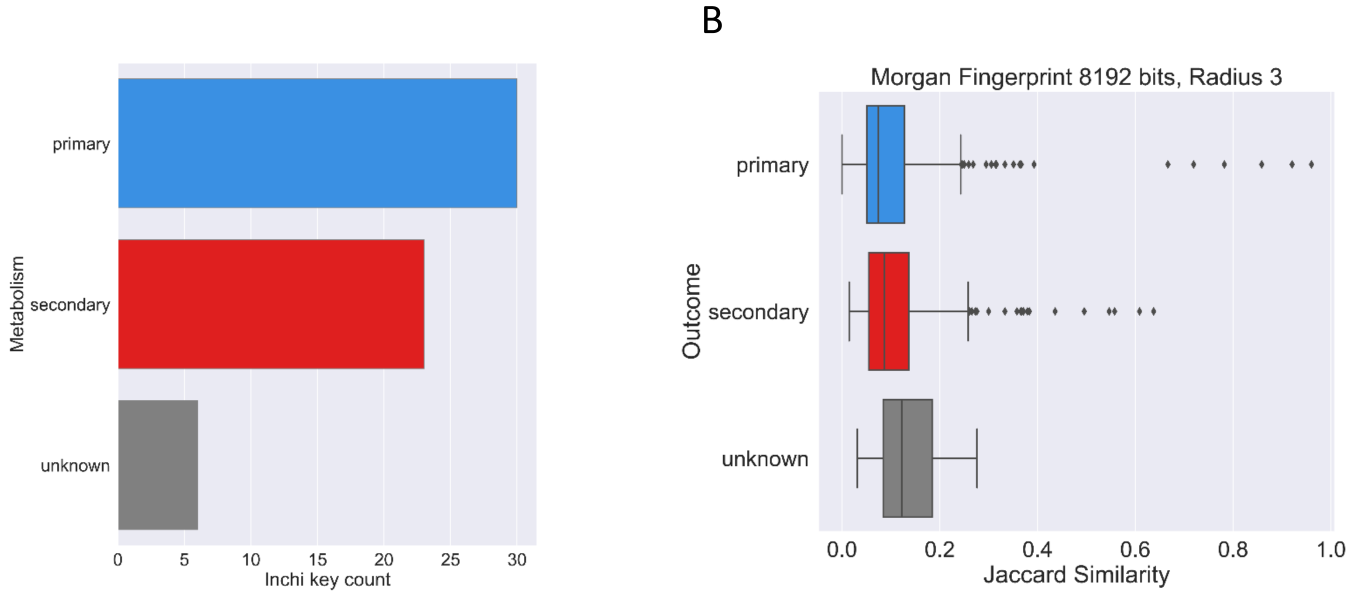


**Figure S6** – Structural similarity analysis of the compound fraction common only to genomics-based annotations and compounds detected experimentally in our lab. (A) Compounds were binned to primary and secondary metabolism according to the pathways they are part of. A secondary classification would be given in cases of a single secondary pathway or multiple pathways where at least one would be secondary. (B) The range of Jaccard similarity as measured pairwise with a 8192 bit vector representation of the Morgan fingerprint.

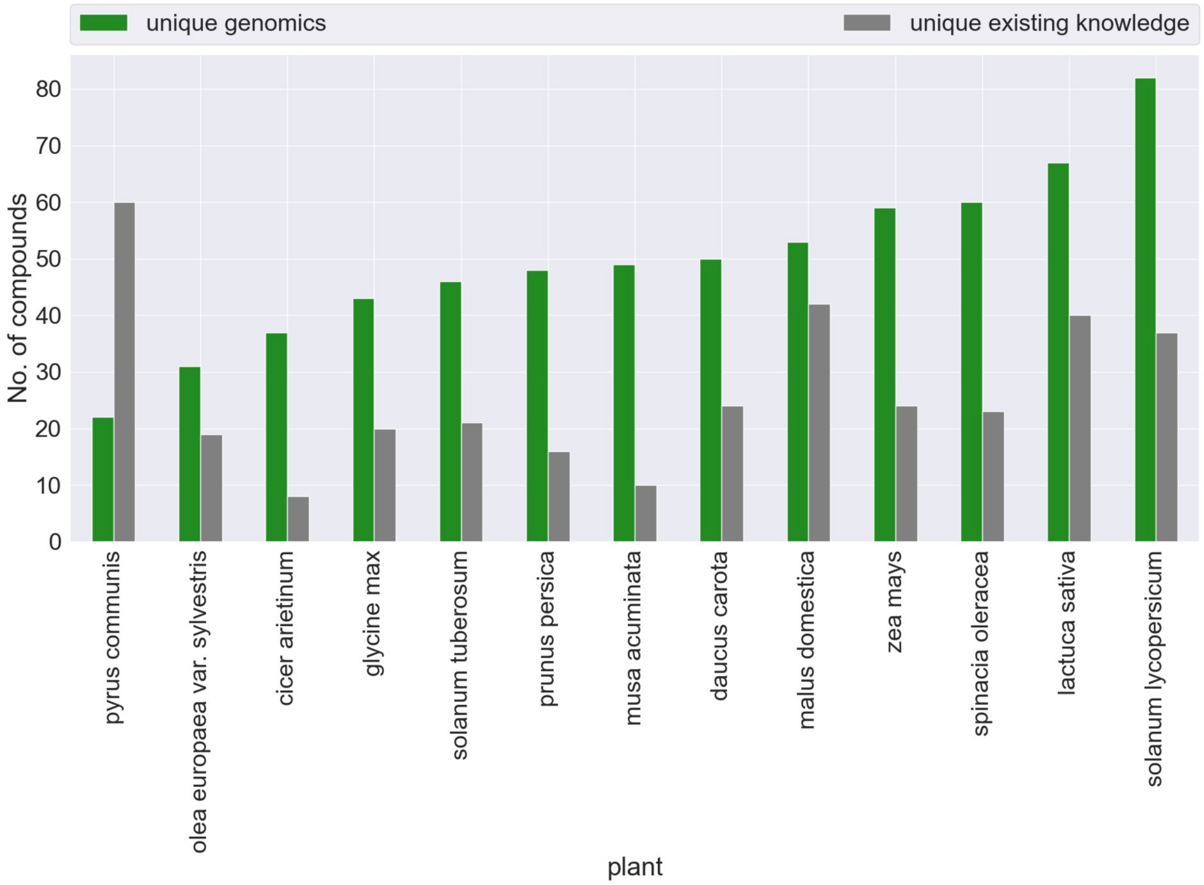


**Figure S7** - a comparison between the unique contribution of compounds (compounds found only in either the genomics-based annotation collection or the existing knowledge collection) for 13 plants in our experimentally detected compound collection.

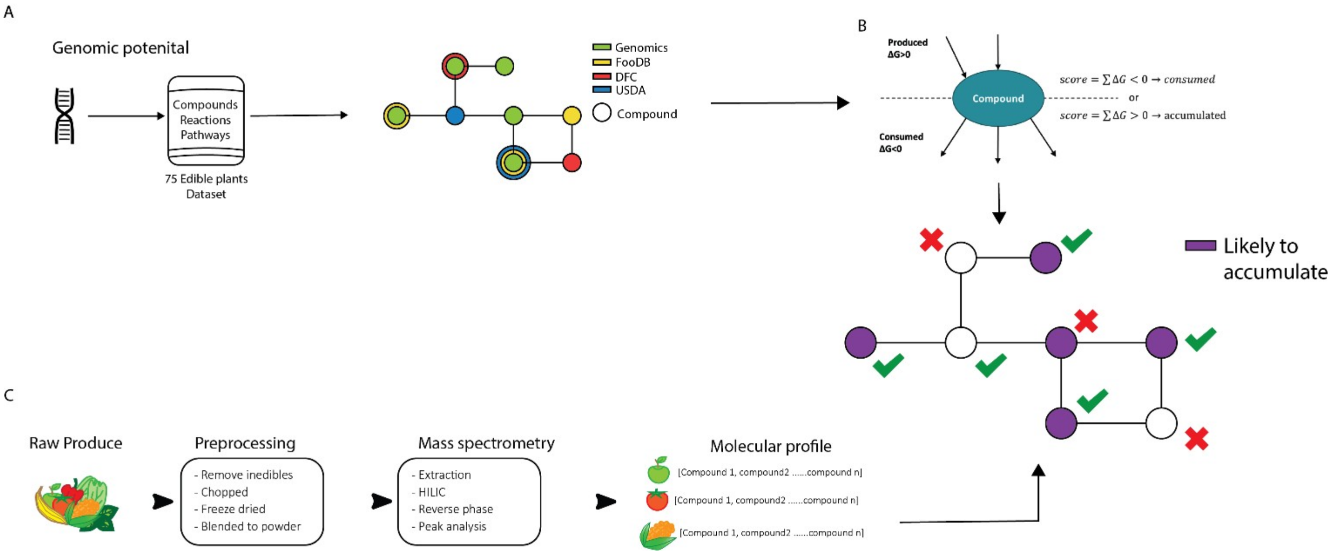


**Figure S8 - A schematic overview of the developed pipeline.** (A) The collection of available genome-associated metabolic annotations and the construction of plant specific genome-scale metabolic networks. (B) A thermodynamics based approach for the prediction of likely to accumulate compounds in plants. Gibbs free energy values were collected from MODElSEED and PlantCyc and a compound score was calculated according to the compounds involvement in all associated reactions (either as a reactant or product). (C) 13 raw produce were sampled from two big chain grocery stores, preprocessed and went into a mass spectrometry analysis to give the molecular profile which was then used as a validation set for the Thermodynamics based approach described in B.

**Table S 1** - Performance metrics for plants included in the thermodynamics feasibility analysis

| plant | Compound fraction | $AUC_{ROC}$ | F-1 score | $AUC_{PR}$ | Optimal threshold | Compound fraction/total scored |
|---|---|---|---|---|---|---|
| cicer arietinum (chickpea) | 210 | 0.767895 | 0.173913 | 0.547619 | 212.4767 | 0.1 |
| daucus carota (carrot) | 310 | 0.761825 | 0.149254 | 0.540323 | 154.2443 | 0.15 |
| glycine max (soybean) | 110 | 0.743802 | 0.181818 | 0.55 | 283.4191 | 0.05 |
| lactuca sativa (lettuce) | 260 | 0.744203 | 0.206897 | 0.557692 | 154.2443 | 0.12 |
| malus domestica (apple) | 260 | 0.721348 | 0.188153 | 0.551923 | 154.2443 | 0.12 |
| musa acuminata (banana) | 210 | 0.741319 | 0.157895 | 0.542857 | 154.2443 | 0.1 |
| olea europaea var. sylvestris (olive) | 160 | 0.689189 | 0.139535 | 0.5375 | 154.2443 | 0.08 |
| prunus persica (peach) | 260 | 0.779375 | 0.142857 | 0.538462 | 154.2443 | 0.12 |
| solanum lycopersicum (tomato) | 210 | 0.762363 | 0.235294 | 0.566667 | 154.2443 | 0.09 |
| solanum tuberosum (potato) | 210 | 0.765776 | 0.165939 | 0.545238 | 154.2443 | 0.09 |
| spinacia oleracea (spinace) | 260 | 0.767094 | 0.181818 | 0.55 | 154.2443 | 0.13 |
| zea mays (corn) | 110 | 0.73614 | 0.24 | 0.568182 | 412.2883 | 0.05 |

**Table S1 legend:**

Compound fraction: the amount of compounds with score above the selected optimal score threshold.

$AUC_{ROC}$ : The area under the curve for the reciever-operator curve.

F-1 score: a standard metric for measuring accuarcy.

$AUC_{PR}$ : The area under the curve for a precision-recall curve.

Optimal threshold: The thermodynamic feasibility score used as threshold as a result of an optimal threshold analysis.

Compound fraction/total scored : the ratio of compounds with scores higher than the threshold and the total number of scored compounds for that plant.
